# A Consensus Model of Glucose-Stimulated Insulin Secretion in the Pancreatic *β*-Cell

**DOI:** 10.1101/2023.03.10.532028

**Authors:** M. Deepa Maheshvare, Soumyendu Raha, Matthias König, Debnath Pal

## Abstract

The pancreas plays a critical role in maintaining glucose homeostasis through the secretion of hormones from the islets of Langerhans. Glucose-stimulated insulin secretion (GSIS) by the pancreatic *β*-cell is the main mechanism for reducing elevated plasma glucose. Here we present a systematic modeling workflow for the development of kinetic pathway models using the Systems Biology Markup Language (SBML). Steps include retrieval of information from databases, curation of experimental and clinical data for model calibration and validation, integration of heterogeneous data including absolute and relative measurements, unit normalization, data normalization, and model annotation. An important factor was the reproducibility and exchangeability of the model, which allowed the use of various existing tools. The workflow was applied to construct the first consensus model of GSIS in the pancreatic *β*-cell based on experimental and clinical data from 39 studies spanning 50 years of pancreatic, islet, and *β*-cell research in humans, rats, mice, and cell lines. The model consists of detailed glycolysis and equations for insulin secretion coupled to cellular energy state (ATP/ADP ratio). Key findings of our work are that in GSIS there is a glucose-dependent increase in almost all intermediates of glycolysis. This increase in glycolytic metabolites is accompanied by an increase in energy metabolites, especially ATP and NADH. One of the few decreasing metabolites is ADP, which, in combination with the increase in ATP, results in a large increase in ATP/ADP ratios in the *β*-cell with increasing glucose. Insulin secretion is dependent on ATP/ADP, resulting in glucose-stimulated insulin secretion. The observed glucose-dependent increase in glycolytic intermediates and the resulting change in ATP/ADP ratios and insulin secretion is a robust phenomenon observed across data sets, experimental systems and species. Model predictions of the glucose-dependent response of glycolytic intermediates and insulin secretion are in good agreement with experimental measurements. Our model predicts that factors affecting ATP consumption, ATP formation, hexokinase, phosphofructokinase, and ATP/ADP-dependent insulin secretion have a major effect on GSIS. In conclusion, we have developed and applied a systematic modeling workflow for pathway models that allowed us to gain insight into key mechanisms in GSIS in the pancreatic *β*-cell.

## 1 INTRODUCTION

The pancreas plays a vital role in maintaining glucose homeostasis (Woods et al., 2006) through the secretion of hormones from the islets of Langerhans. The most important hormones are insulin, secreted by the pancreatic *β*-cells, and glucagon, secreted by the *α*-cells, both of which play key roles in regulating glucose homeostasis (König et al., 2012a).

Glucose-induced insulin secretion (GSIS) is a physiological process by which the pancreas releases insulin in response to an increase in blood glucose levels. When glucose enters the bloodstream after a meal, it is taken up by *β*-cells in the pancreas through glucose transporters, primarily GLUT2 (MacDonald et al., 2005). Once inside the *β*-cells, glucose is metabolized via glycolysis, which produces energy in the form of ATP.

The coupling of glycolysis with the insulin secretion mechanism in the *β*-cell is established by the regulatory effects of glycolytic intermediates on the levels of energy metabolites such as ATP and NADH (Newsholme et al., 2014; Prentki et al., 2013). The rise in ATP levels triggers a series of events that lead to the release of insulin. Specifically, the high ATP levels close ATP-sensitive potassium channels (Ashcroft, 2006), which leads to depolarization of the cell membrane and opening of voltage-gated calcium channels. The influx of calcium triggers the exocytosis of insulin-containing vesicles, leading to the release of insulin into the bloodstream (Rorsman and Braun, 2013; Guerrero-Hernandez and Verkhratsky, 2014). The *K*_*AT P*_ /*Ca*^2+^ independent signaling mechanisms and the other metabolites besides glucose contribute to the amplification of the signaling events that trigger insulin secretion (Guay et al., 2013).

GSIS by the pancreatic *β*-cell is the primary mechanism for lowering elevated plasma glucose levels. The amount of insulin released increases with the glucose in the bloodstream. This process is crucial for the regulation of blood glucose levels by promoting the uptake and use of glucose by cells throughout the body, such as muscle, fat tissue, and the liver (Di Camillo et al., 2014; Fritsche et al., 2008).

Glycolysis is the primary metabolic pathway responsible for GSIS. It involves the uptake of glucose and its conversion to pyruvate, which is critical for ATP synthesis and maintenance of ATP levels. Experimental data from metabolic profiling studies in islet cells support the key role of glycolysis in GSIS (Spégel et al., 2013, 2015; Taniguchi et al., 2000). As glucose levels increase, glycolytic flux and most glycolytic intermediates increase in a dose-dependent manner. Changes in adenine nucleotide levels due to variations in glycolytic flux lead to changes in nucleotide ratios, with increasing glucose levels resulting in a positive correlation between the ATP/ADP ratio and Ca^2+^ response and insulin release. This trend is consistent across several studies (Detimary et al., 1996; Malaisse et al., 1978; Salvucci et al., 2013), including isolated islets perfused with glucose, rat and mouse tissue homogenates, and insulin-secreting cell lines. The increase in ATP/ADP ratio ranges from 2 to 7 when glucose levels are increased from 2.8mM to 30mM, indicating similar behavior in different experimental systems studying insulin secretion by the pancreas (Huang and Joseph, 2014).

Mathematical models have been developed to investigate the metabolic and signaling mechanisms that trigger and amplify insulin secretion. Early models of *β*-cells focused on examining the relationship between glycolytic oscillations and pulsatile insulin release to understand GSIS (Bertram et al., 2007; Tornheim, 1997). Merrins et al. analyzed the oscillations in glycolytic intermediates (i.e. fructose-6-phosphate, fructose-2,6-bisphosphate, and fructose-1,6-bisphosphate) and their effect on pulsatile insulin secretion (Merrins et al., 2012), while other models integrated glycolytic flux with mitochondrial ATP production to study the role of reducing equivalents such as pyridine nucleotides in enhancing insulin secretion (Westermark et al., 2007; Bertram et al., 2006). Jiang et al. further combined previously developed models of glycolysis, citric acid cycle, *β*-oxidation, pentose phosphate shunt, and respiratory chain and studied the local and global dynamics of the GSIS mechanism in response to parameter perturbations. These models were coupled with the calcium signaling pathway of Fridyland et al. to create an integrated metabolic model (Fridlyand and Philipson, 2010; McKenna et al., 2016).

To investigate the synergistic insulinotropic effect of other nutrient sources, Salvucci et al. (Salvucci et al., 2013) developed a model by integrating alanine metabolism with glucose metabolism, the citric acid cycle, and the respiratory chain. Gelbach et al. developed a system of 65 reactions integrating glycolysis, glutaminolysis, the pentose phosphate pathway, the citric acid cycle, the polyol pathway, and the electron transport chain to study the kinetics of insulin secretion (Gelbach et al., 2022).

However, the majority of these models are based on earlier models that were developed using kinetic data from organisms other then humans or non-pancreatic tissues, such as a glycolysis model that utilized kinetic data from experiments on yeast cell extract, or a glycolysis model based on kinetic data from mammalian muscle (Smolen, 1995). Often, the data used to build these models is limited and comes from a single experimental study. In most models specific to *β*-cells, reaction kinetics are described by simple mass-action rate laws. There exists no detailed kinetic model of the changes in glycolysis during GSIS that can effectively integrate the observed changes in glycolytic and energy intermediates from a wide range of GSIS experiments.

In systems biology and systems medicine, ensuring the reproducibility of computational models and integrating diverse data from multiple sources into these models are critical challenges. Standards for model description, such as the Systems Biology Markup Language (SBML) (Hucka et al., 2015; Keating et al., 2020), have been developed to enable the reusability and reproducibility of existing models, but they have yet to be utilized in the field of pancreatic GSIS modeling. Furthermore, there is a need to address how to integrate heterogeneous data from different studies conducted in different organisms and experimental systems in the context of GSIS modeling.

This study aims to develop a detailed kinetic model of GSIS and the associated changes in glycolysis in the pancreatic *β*-cell. The novel contributions of this work include a systematic curation and integration of changes in glycolytic metabolites from different experimental studies across different species and experimental systems. Based on this unique data set, a detailed kinetic model of glycolysis and GSIS was constructed using a systematic approach with a focus on reproducibility. This approach allowed the establishment of a consensus model of the changes that occur in insulin secretion with varying glucose concentrations. The overall goal was to provide a better understanding of the mechanisms underlying GSIS and to contribute to the development of improved computational models of these processes.

## 2 RESULTS

Our study introduces a detailed kinetic model of GSIS in the pancreatic *β*-cell, which has the ability to simulate alterations in glycolytic intermediates and ATP/ADP ratio due to glucose levels and the effect of change in the energy state of the *β*-cell on insulin secretion.

### 2.1 Systematic curation of data set of changes in GSIS

In the course of this study, we compiled a comprehensive data set (Tab. 1) of GSIS based on experimental and clinical data from 39 studies spanning half a century of research on pancreatic, islet, and *β*-cell function in humans, rats, mice, and cell lines. Specifically, we systematically curated metabolomics data from studies conducted between 1970 and 2020, comprising information on the concentration of glycolytic intermediates and cofactors in both time-course and steady-state experiments, as well as the corresponding glucose doses. The data set contains 17 metabolites, comprising 359 data points from steady-state experiments and 249 data points from time-course studies. It includes both absolute and relative measurements of metabolite changes, and an overview of the available information for each metabolite and study is presented in Fig. 1.

**Figure 1.**
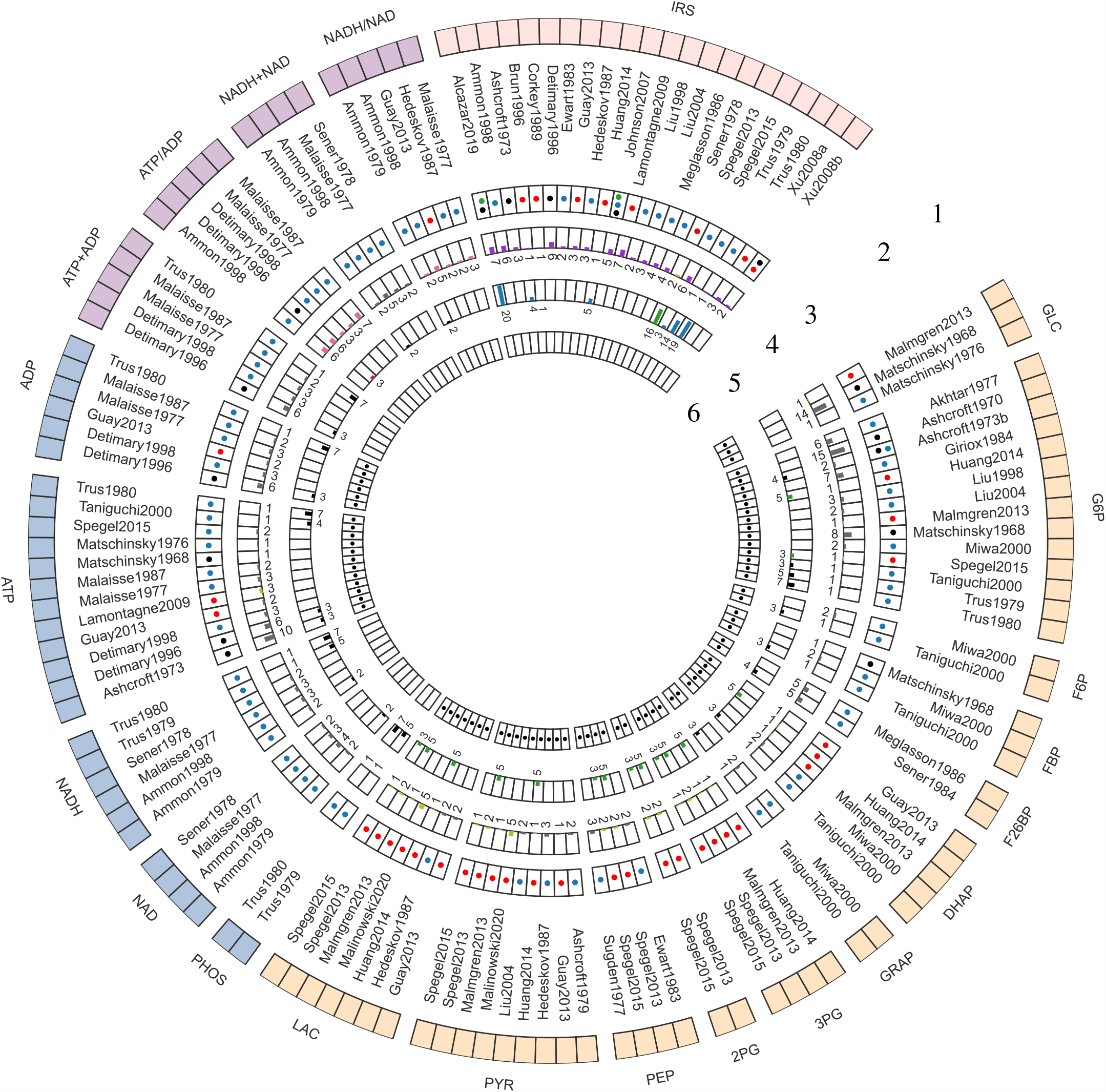
Curated data for model development and evaluation. The data description is detailed from the periphery to the center of the Circos plot. *1. Model elements:* The outermost layer provides an overview of the metabolites included in the data set. GLC: glucose, G6P: glucose 6-phosphate, F6P: fructose 6-phosphate, FBP: fructose 1,6-bisphosphate, F26BP: fructose 2,6-bisphosphate, DHAP: dihydroxyacetone phosphate, GRAP: glyceraldehyde 3-phosphate, BPG: 1,3-biphosphoglycerate, 3PG: 3-phosphoglycerate, 2PG: 2-phosphoglycerate, PEP: phosphoenolpyruvate, PYR: pyruvate, LAC: lactate, PHOS: phosphate, NAD: nicotinamide adenine dinucleotide, NADH: reduced nicotinamide adenine dinucleotide, NADH total: NADH + NAD; NADH ratio: NADH/NAD; ATP: adenosine triphosphate, ADP: adenosine diphosphate, ATP total: ATP + ADP, ATP ratio: ATP/ADP, IRS: insulin secretion rate. The metabolites were grouped in the following categories: Color code: 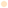 glycolytic intermediates, 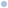 cofactors, 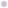 cofactor ratio or sum, 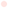 insulin secretion rate (IRS); *2. Studies:* The second layer depicts the islet-cell specific metabolite profiling studies curated from the literature; *3. Animal species:* The third layer indicates the animal species or cell line from which the data was curated. Color code: 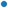 Rat, 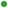 Human, • Mouse, and 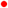 Cell line data; *4. time course data:* The fourth layer shows a bar graph illustrating the number of data points collected from studies reporting time course data of metabolites. Color code: 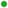 relative (or fold), • concentration, 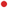 ratio, 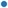 rate measurements; *5. Steady-state data:* The fifth layer indicates the number of data points collected from studies reporting steady-state/ dose-response data of metabolites. Color code: 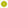 relative (or fold), 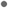 concentration, 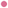 ratio, 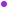 rate measurements; *6. Data used for parameter estimation*: The innermost layer indicates the subset of data used for parameter fitting.

This data set represents the first open and FAIR (findable, accessible, interoperable, and reusable) large-scale collection of data on changes in glycolysis and insulin secretion in the pancreatic *β*-cell during GSIS. We used the absolute and relative measurements of glycolysis metabolites and insulin secretion rates in this data set for model calibration and evaluation.

The data set is available under a CC-BY4.0 license from https://github.com/matthiaskoenig/pancreas-model.

### 2.2 Reproducible modeling workflow

In this study, we describe a comprehensive modeling workflow for building small kinetic pathway models (Fig. 2) using SBML (Hucka et al., 2015; Keating et al., 2020).

**Figure 2.**
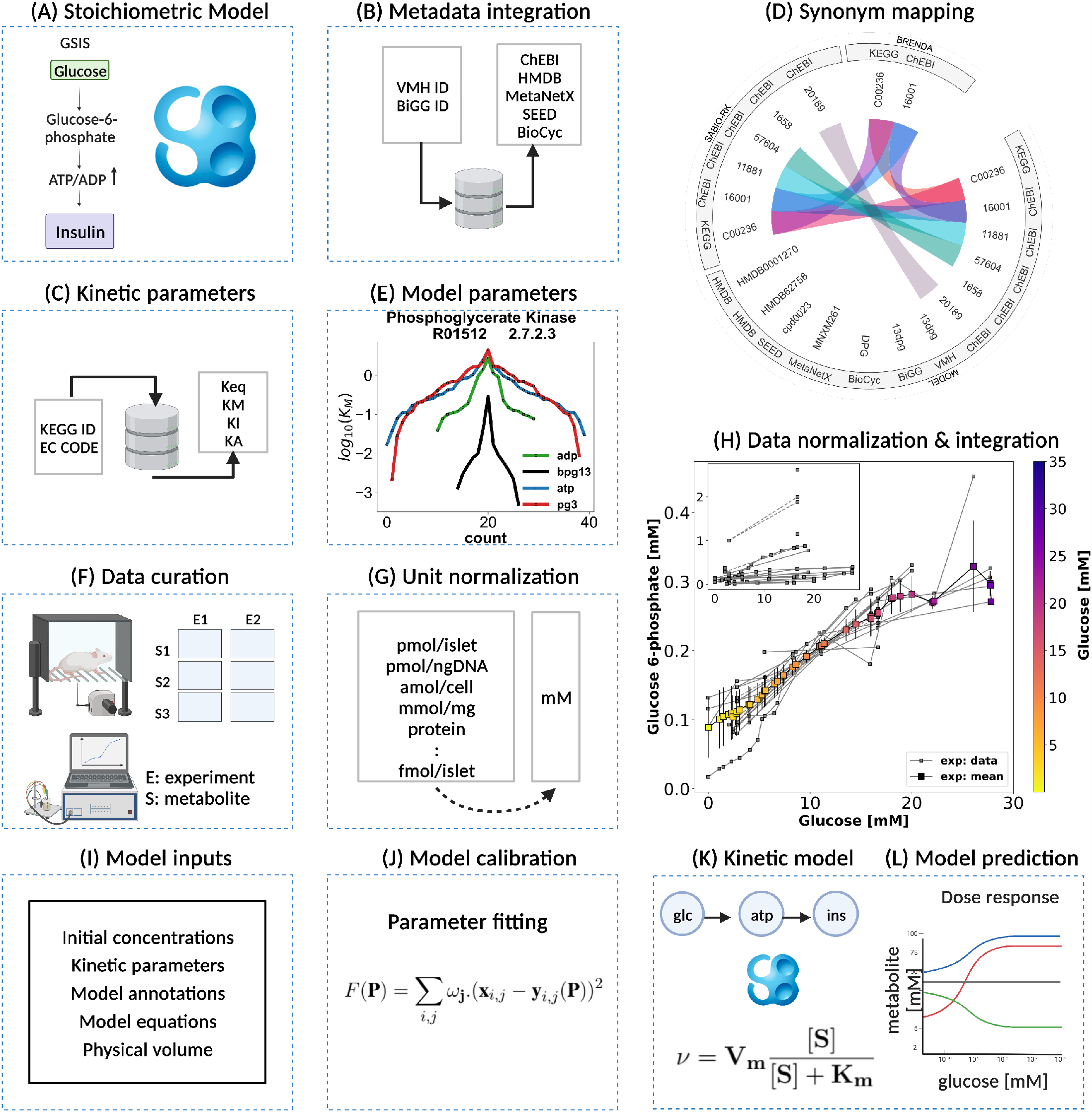
Kinetic model development workflow. *(A) Initial stoichiometric model in SBML*. Glycolytic reactions were collected from VMH database and existing models of glycolysis. *(B) Metadata integration*. VMH and BiGG database field identifiers were used to retrieve additional metadata such as HMDB, BioCyc, MetaNetX, ChEBI, and SEED database field identifiers. *(C) Synonym mapping*. The synonyms associated with each metabolite were queried using compound identifier mapping services. *(D) Kinetic parameters*. EC number and KEGG reaction identifiers were used to query half-saturation/Michaelis-Menten *K*_*M*_, inhibition *K*_*I*_, activation *K*_*A*_, and equilibrium *K*_*eq*_ constants (synonym mapping was applied for all compounds). *(E) Model parameters*. The parameter values retrieved from different databases were merged and median values were assigned to the model parameters; *(F) Data curation*. A systematic literature search was performed and metabolite concentrations from islet cell studies were curated. *(G) Unit normalization*. Absolute and relative quantification of metabolite concentrations reported in heterogeneous units were converted to mM. *(H) Data normalization*. Systematic bias observed in the unit-normalized data was removed by performing least-squares minimization to minimize the distance between the mean curve of the unit-normalized data curves and the experimental curves of the unit-normalized data. *(I) Model inputs*. Values of kinetic parameters, initial concentrations, volumes, equations, and annotations have been assigned to the model entities. *(J) Model calibration*. Time course and steady-state data were used for parameter estimation. *(K) Kinetic SBML model*. The final kinetic SBML model was generated. *(L) Model prediction*. Glycolytic intermediates and insulin response were predicted as a function of varying glucose concentrations. Created with BioRender.com.

In our model-building workflow, we followed several steps to construct a kinetic SBML model of glycolysis. A) First, we built an SBML model based on glycolytic reactions and intermediates from existing models and pathway databases. B) We then annotated metabolites and reactions with metadata information which was extended by querying VMH and the BiGG database, resulting in mappings to additional resources such as HMDB, BioCyc, MetaNetX, ChEBI, and SEED. C) We collected and retrieved kinetic parameters such as *K*_*M*_, *K*_*I*_, *K*_*A*_, and *K*_*eq*_ constants from databases and D) integrated them with synonyms associated with each queried metabolite using compound identifier mapping services. E) We integrated the resulting parameters and assigned median values to the model parameters. F) Next, we curated data from studies reporting metabolite concentrations and changes, and insulin secretion in pancreatic, islet, and *β*-cell lines through a literature search. G) Unit normalization was then performed to convert reported metabolite concentrations and insulin secretion to mmole/l (mM) and nmole/min/ml (*β*-cell volume), respectively. H) Data normalization was performed to remove systematic differences between data reported in different studies and experimental systems. I) Next, values for kinetic parameters, initial concentrations, volumes, rate equations, and annotations were integrated into the stoichiometric model. J) We calibrated the model by parameter optimization using time-course and steady-state data and K) generated the final SBML kinetic model using all the information. L) Finally, we performed model predictions of glycolytic intermediates and insulin response as a function of varying glucose concentrations. Steps were performed iteratively to fill gaps and extend the data set and model.

### 2.3 Computational model

Using the established data set, we utilized the aforementioned workflow to develop the first consensus model of GSIS in the pancreatic *β*-cell. The model is comprised of detailed glycolysis and equations for insulin secretion which are coupled to the cellular energy state (ATP/ADP ratio). The metabolites and reactions incorporated into the kinetic model are depicted in Fig. 3, and their biochemical interactions are represented through a system of ordinary differential equations. The model consists of 21 enzyme-catalyzed reactions, 25 metabolites, and 91 parameters, and also includes an empirical model that connects the energy state of the *β*-cell to insulin secretion.

**Figure 3.**
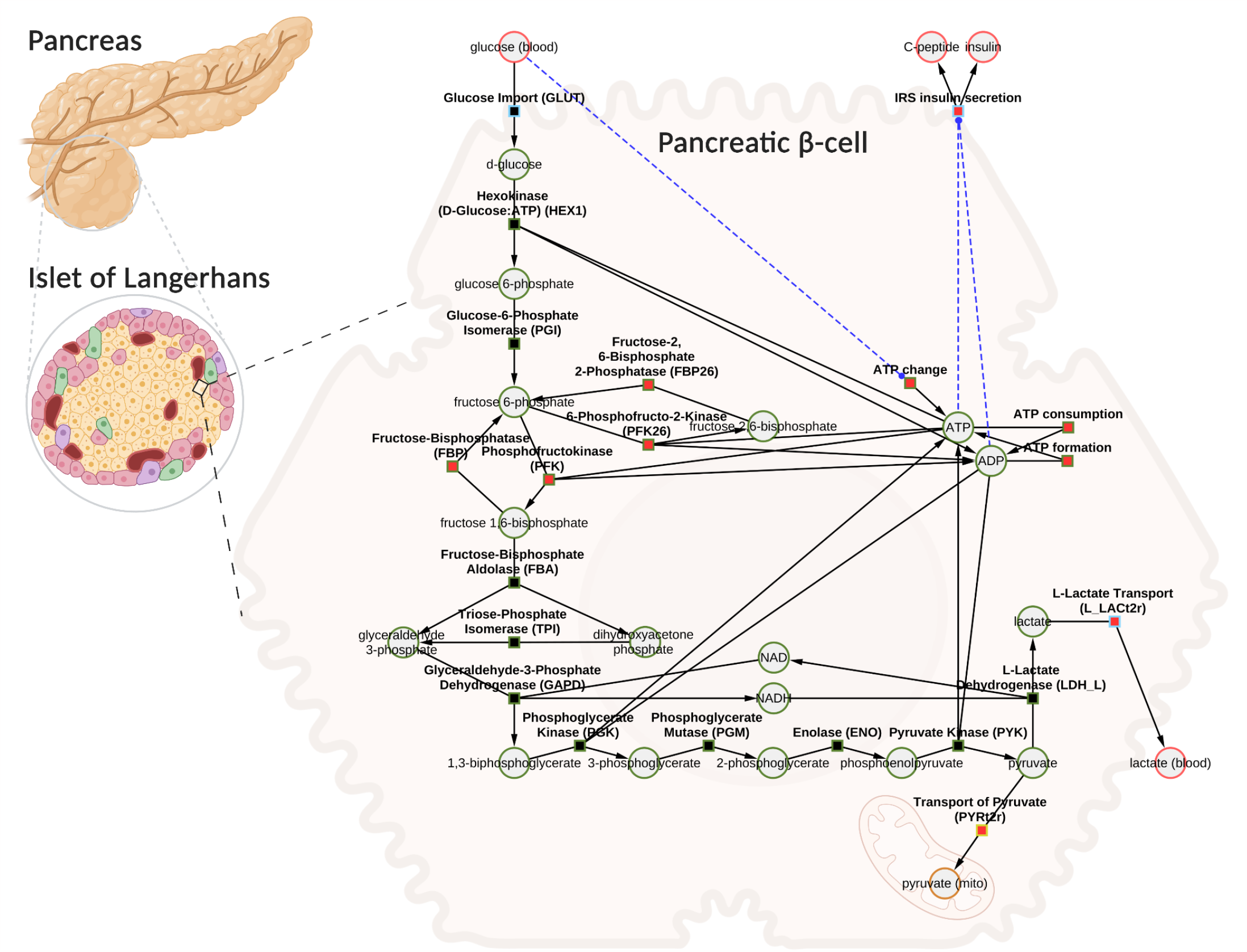
Computational model of glucose-stimulated insulin secretion (GSIS) in the pancreatic *β*-cell. The model consists of glycolysis and insulin secretion coupled to the energy state (ATP/ADP ratio). The GLUT transporter facilitates the uptake of glucose from the plasma into the cell. Glucose undergoes phosphorylation and the subsequent reactions lead to the production of pyruvate. Pyruvate can either be converted to lactate and exported into blood or transported to the mitochondria where it serves as a fuel source for the production of tricarboxylic acid cycle (TCA) intermediates (the TCA cycle has not been modeled). Depending on the external glucose concentrations, glycolysis intermediates and energy metabolites such as ATP, ADP, NAD, and NADH change. An increase in the ATP/ADP ratio as a result of changes in glucose triggers the cascade of signaling mechanisms that promote insulin secretion by the pancreatic *β*-cell. Phosphate, water, and hydrogen ions have been omitted from the diagram for clarity (but are included in the model for mass and charge balance). The network diagram was created using CySBML (König et al., 2012b). Created with BioRender.com.

When glucose levels are high, GLUT transporter allows glucose to enter the cell, and glucokinase converts glucose to glucose-6-phosphate. The upper glycolysis produces fructose-6-phosphate, fructose-1,6-phosphate, and triose phosphates like dihydroxyacetone phosphate and glyceraldehyde phosphate. Lower glycolysis then leads to the creation of 3-phosphoglycerate, 2-phosphoglycerate, phosphoenolpyruvate, and pyruvate. Pyruvate can be transformed into lactate or transported to the mitochondria. For each glucose molecule, two ATP molecules are produced. Changes in ATP/ADP ratio trigger insulin secretion.

The SBML model is available under a CC-BY4.0 license from https://github.com/matthiaskoenig/pancreas-model.

### 2.4 Normalization of data

The aim of this study was to investigate variations in glycolysis, glycolytic intermediates, energy metabolites, and insulin secretion during GSIS using the established model. In order to integrate heterogeneous experimental data for each metabolite and insulin secretion rate, we conducted a two-step normalization process to standardize time course and dose-response measurements. The normalization process involved unit normalization (as discussed in Sec. 4.7) and data normalization (as discussed in Sec. 4.8) to normalize the diverse data and eliminate systematic deviations for individual studies. We present the case of glucose 6-phosphate as an example of the normalization process (see Fig. 4). The experimental curves were converted to relative (fold) and unit-normalized absolute measurements (Fig. 4A and Fig. 4B). To combine the fold data and absolute data, we multiplied the fold values by the basal concentration to obtain absolute values (Fig. 4C). If the basal metabolite concentration was not reported, we used the mean curve of the absolute data at the pre-incubation glucose dose of the experiment to determine the basal value. For metabolites consisting of only relative measurements, we used the half-saturation *K*_*m*_ value of the metabolite as an estimate for the basal concentration. Using this strategy, we converted all fold-changes and time courses to absolute data with standardized units, which was then combined with the existing absolute data. However, the standard deviation of the combined data set measurements was high, and large systematic differences between studies could be observed. We determined scaling factors for every study to minimize the difference between all studies based on least-squares minimization (as discussed in Sec. 4.8.1). The resulting normalized data (Fig. 4D) was then used for model calibration. We applied this procedure to all metabolites in the model as well as the insulin secretion rate, reducing the variability in the data substantially.

**Figure 4.**
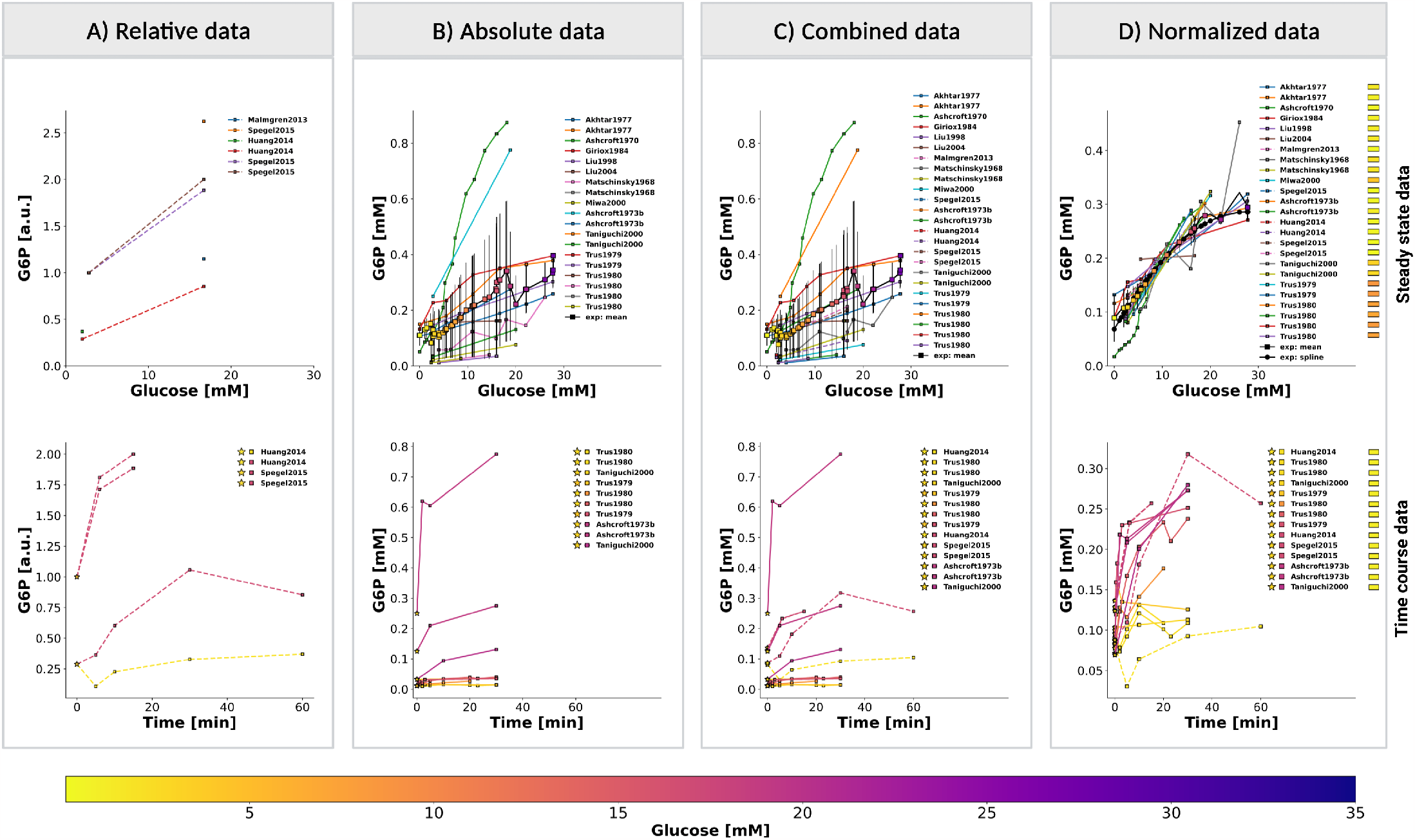
Normalization of steady-state and time course data for glucose 6-phosphate (G6P). ***(A)*** *Relative data*. Experimental curves from *β*-cell studies reporting relative levels of G6P, expressed as fold to baseline value; ***(B)*** *Absolute data*. Experimental curves from *β*-cell studies reporting absolute concentrations of G6P, the plot displays the unit-normalized absolute data. ***(C)*** *Combined data*. The relative (fold) measurements were converted to absolute units and combined with the unit-normalized absolute data. ***(D)*** *Normalized data*. Systematic biases between different studies of the combined data were removed by data normalization. Data normalization was performed by minimizing the offset (sum of squared residuals) between the mean curve and the experimental curves. The *mean curve* was computed as the weighted average of the experimental curves and *spline curve* is the piecewise-polynomial interpolation of the data points in the mean curve. For steady-state data, the legend indicates studies associated with the experimental curves. For time course data, the legend indicates the pre-incubation glucose dose (⭐), incubation glucose dose (□), experimental study, and the value of scale transformation parameter *f* ^*α*^ (▭) of experiment *α. (top panel)* and *(bottom panel)* show the data of dose-response and time course experiments, respectively. Data from (Akhtar et al., 1977; Ashcroft et al., 1970, 1973b; Giroix et al., 1984; Huang and Joseph, 2014; Liu et al., 1998, 2004; Malmgren et al., 2013; Matschinsky and Ellerman, 1968; Miwa et al., 2000; Spégel et al., 2015; Taniguchi et al., 2000; Trus et al., 1979, 1980). For more details, please refer to Sec. 2.1.

### 2.5 Changes in glycolytic metabolites and insulin secretion in GSIS

Our work has uncovered several key findings related to GSIS. First, we found that almost all glycolytic intermediates increase in a glucose-dependent manner across a wide range of glucose concentrations, as illustrated in Figures 5, 6, and 7. This increase in glycolytic intermediates is accompanied by a corresponding increase in energy metabolites, especially ATP and NADH. However, one notable exception is ADP, which decreases with increasing glucose levels. As a result, there is a significant increase in ATP/ADP ratios in *β*-cells with increasing glucose, a key factor in insulin secretion. This phenomenon is robust across different data sets, experimental systems, and species. An important observation is that not only ATP and NADH increase with increasing glucose, but also the total ATP (ATP + ADP) and total NADH (NAD + NADH).

**Figure 5.**
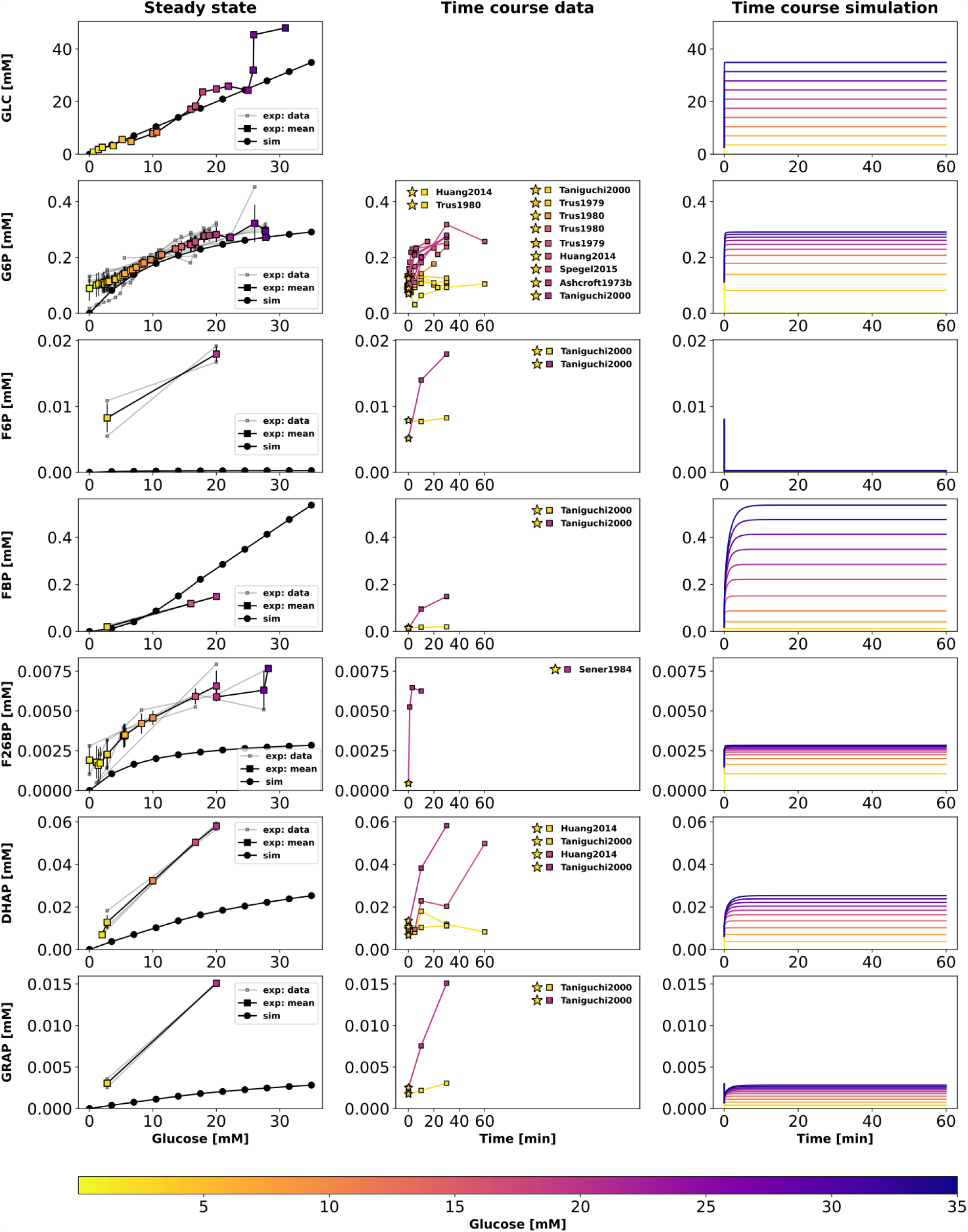
Effect of variations in blood glucose on glycolytic intermediates. *(left column) Dose-response simulations*. Glucose scan was performed for the calculation of steady-state concentration of metabolites in the model. The steady-state concentrations predicted by the model at various glucose doses were compared with the normalized values of experimental measurements; *(middle column) Time course experimental data*. Time course values of glycolytic intermediates and cofactors from multiple experimental studies carried out at different incubation doses of glucose; (⭐) in the legend indicates the pre-incubation glucose dose. *(right column) Time course simulations*. The effect of variation in blood glucose dose on the transient concentration of metabolites. GLC: glucose, G6P: glucose 6-phosphate, F6P: fructose 6-phosphate, FBP: fructose 1,6-bisphosphate, F26BP: fructose 2,6-bisphosphate, DHAP: dihydroxyacetone phosphate, GRAP: glyceraldehyde 3-phosphate. Data from (Akhtar et al., 1977; Ashcroft et al., 1970, 1973b; Giroix et al., 1984; Huang and Joseph, 2014; Liu et al., 1998, 2004; Malmgren et al., 2013; Matschinsky and Ellerman, 1968; Miwa et al., 2000; Spégel et al., 2015; Sener et al., 1984; Taniguchi et al., 2000; Trus et al., 1979, 1980).

**Figure 6.**
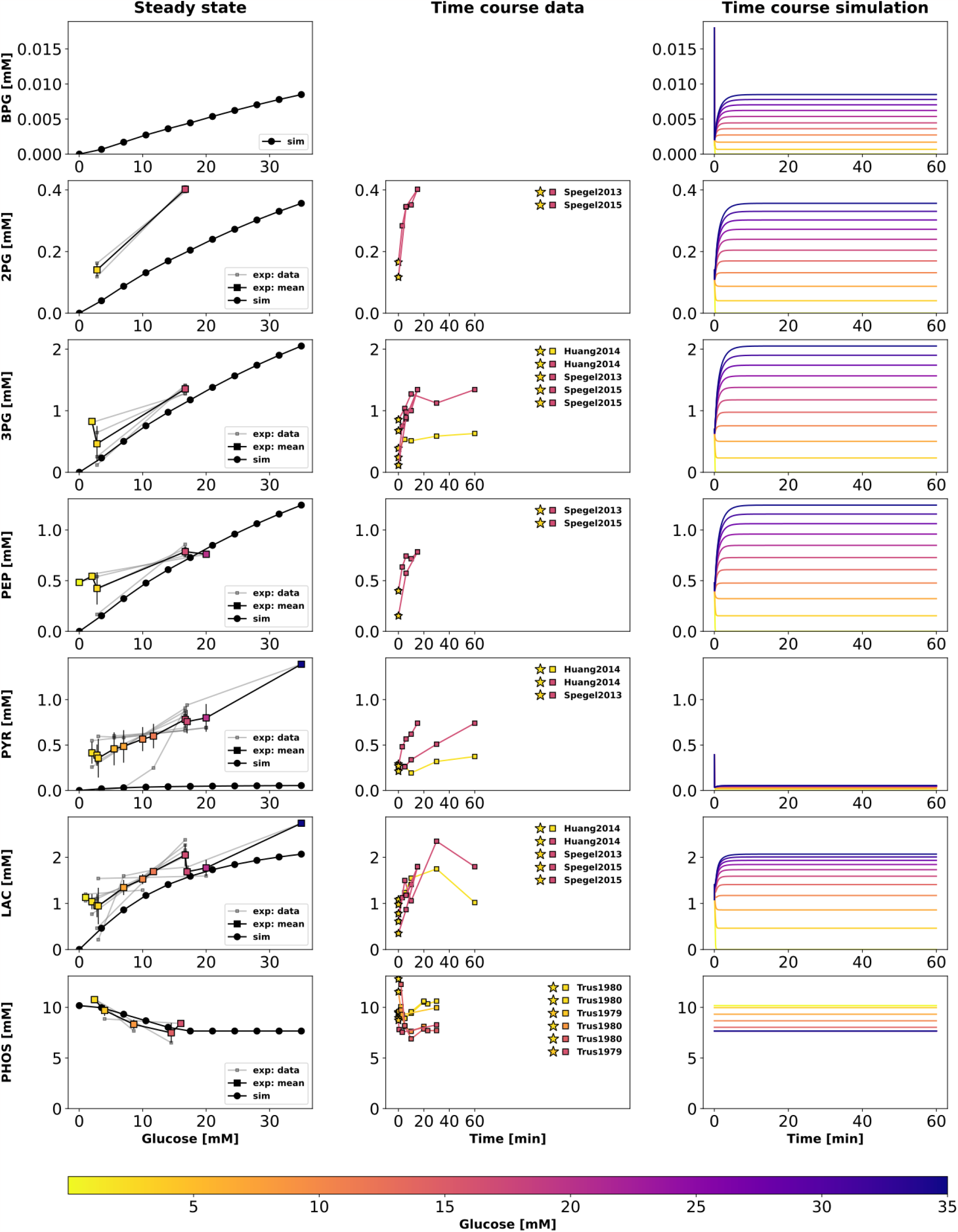
Effect of variations in blood glucose on glycolytic intermediates. The plot is analogous to Fig. 5. BPG: 1,3-biphosphoglycerate, 2PG: 2-phosphoglycerate, 3PG: 3-phosphoglycerate, PEP: phosphoenolpyruvate, PYR: pyruvate, LAC: lactate, PHOS: phosphate. Data from (Ashcroft and Christie, 1979; Ewart et al., 1983; Guay et al., 2013; Hedeskov et al., 1987; Huang and Joseph, 2014; Malinowski et al., 2020; Malmgren et al., 2013; Spégel et al., 2013, 2015; Sugden and Ashcroft, 1977; Trus et al., 1979, 1980).

**Figure 7.**
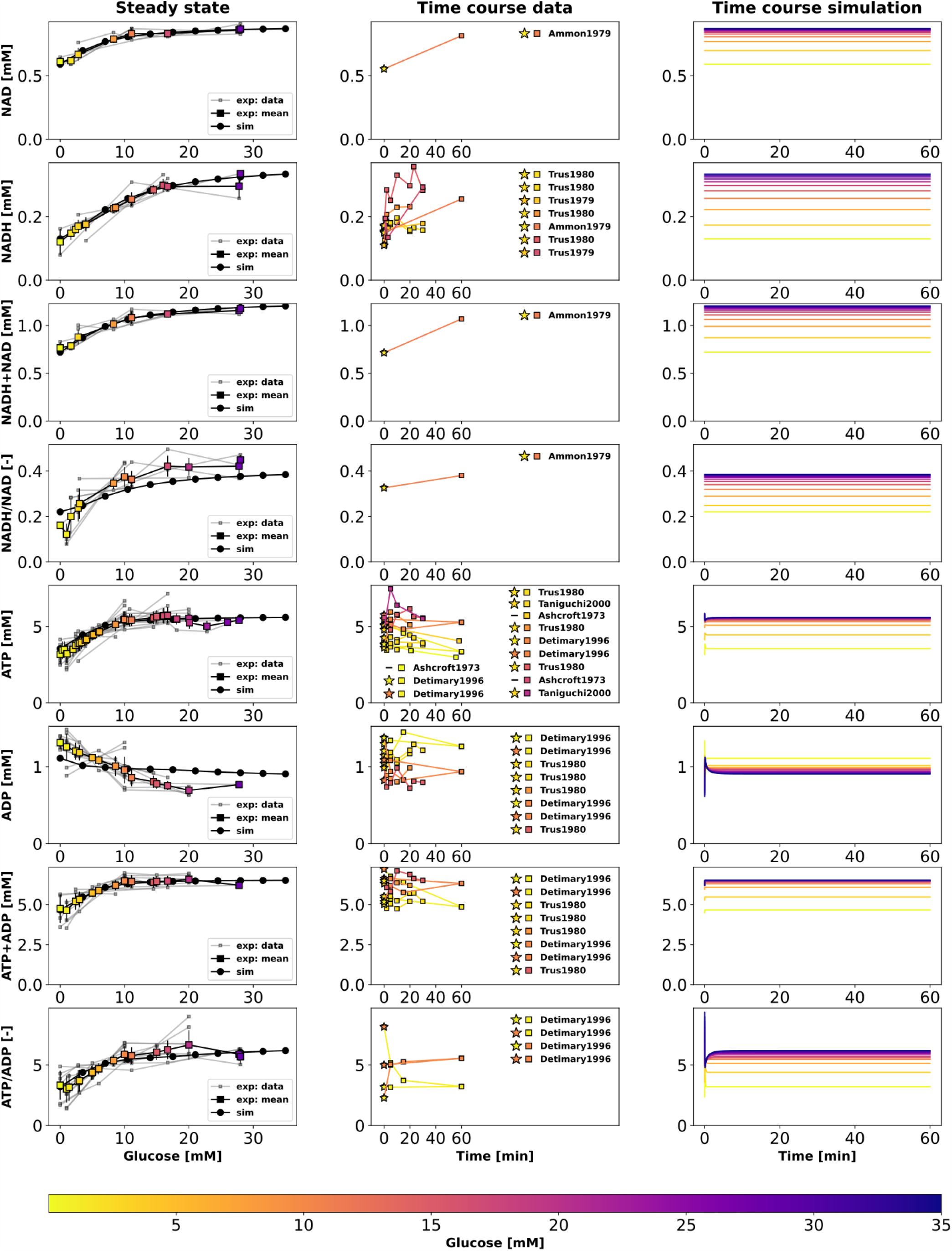
Effect of variations in blood glucose on glycolytic cofactors. The plot is analogous to Fig. 5. NAD: nicotinamide adenine dinucleotide, NADH: nicotinamide adenine dinucleotide reduced. ATP: adenosine triphosphate, ADP: adenosine diphosphate Data from (Ammon et al., 1979, 1998; Ashcroft et al., 1970; Detimary et al., 1996, 1998; Guay et al., 2013; Hedeskov et al., 1987; Lamontagne et al., 2009; Malaisse et al., 1978; Malaisse and Sener, 1987; Matschinsky and Ellerman, 1968; Matschinsky et al., 1976; Sener et al., 1978; Spégel et al., 2015; Taniguchi et al., 2000; Trus et al., 1979, 1980).

Our model was able to predict the glucose-dependent response of glycolytic intermediates and insulin secretion with good agreement to most experimental measurements, as summarized in Table 1. We observed a dose-dependent increase in glycolytic intermediates when glucose concentrations were increased from 0.01 mM to 35 mM. The model predicts that steady states of glycolytic metabolites under constant glucose are reached after approximately 20 minutes, with only 5-10 minutes required to reach steady state according to our simulations.

**Table 1.**
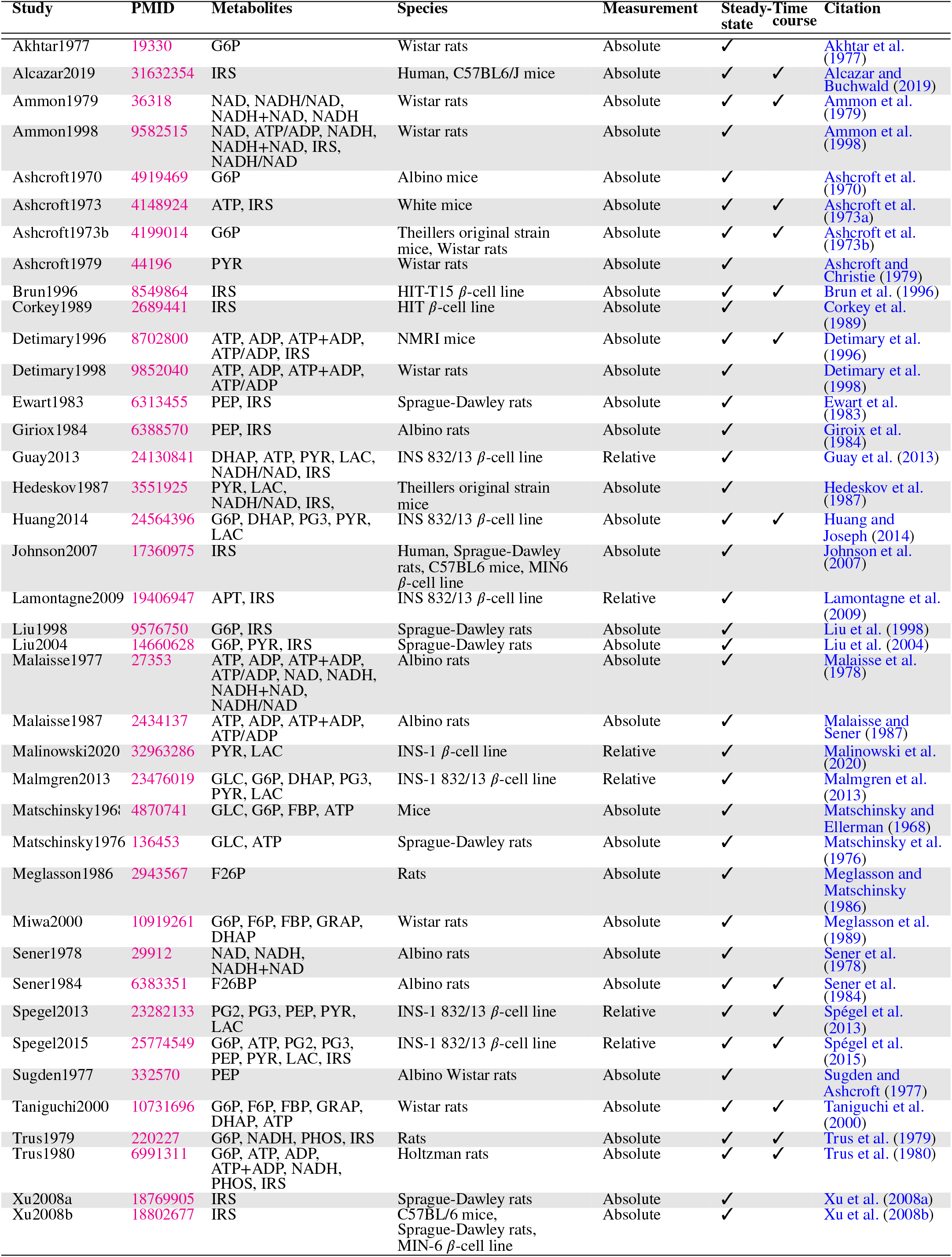
Overview of studies reporting concentrations of metabolites used for model calibration.

Figure 8A illustrates the relationship between glucose dose and insulin release, while Figure 8B shows the effect of varying the ATP/ADP ratio on the insulin response. Specifically, the ATP and ADP concentrations of the *β*-cell increase and decrease, respectively, with the external glucose dose, resulting in an increased ATP/ADP ratio that triggers insulin release. The model is able to reproduce the steady-state insulin secretion depending on glucose concentration, but fails to describe the fast initial insulin release.

**Figure 8.**
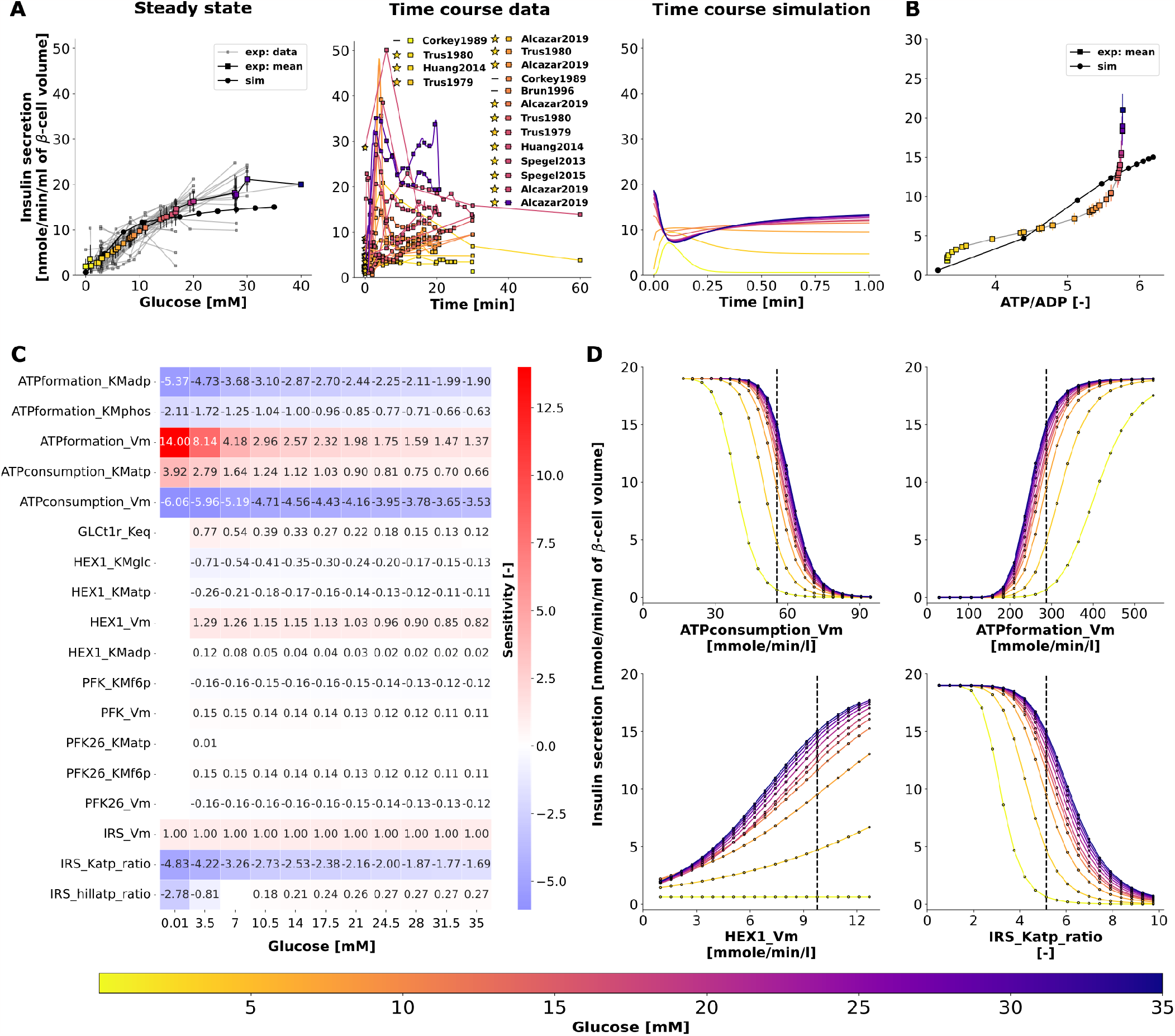
(A) Effect of variations in blood glucose on insulin secretion. The plot is analogous to Fig. 5. Data from (Alcazar and Buchwald, 2019; Ammon et al., 1998; Ashcroft et al., 1973a; Brun et al., 1996; Corkey et al., 1989; Detimary et al., 1996, 1998; Ewart et al., 1983; Guay et al., 2013; Hedeskov et al., 1987; Huang and Joseph, 2014; Johnson et al., 2007; Lamontagne et al., 2009; Liu et al., 1998, 2004; Meglasson and Matschinsky, 1986; Sener et al., 1978; Spégel et al., 2013, 2015; Trus et al., 1979, 1980; Xu et al., 2008a,b). **(B) Effect of change in energy state (ATP/ADP ratio) of the** *β***-cell on insulin secretion**. The rate of insulin release in response to changes in ATP/ADP ratio is shown. **(C) Sensitivity analysis indicating the effect of perturbation in model parameters on insulin secretion**. Heatmap illustrating the values of scaled local sensitivities illustrating the effect of parameter perturbations on the amount of insulin secretion at varying glucose doses. Highly sensitive values are colored in red and blue. The parameters which cause less than 1% change in insulin response for 10% perturbation were not displayed for clarity. For more details, please refer to Sec. 2.6. **(D) Effect of change in model parameters on insulin secretion as a function of glucose dose**. The rate of insulin secretion in response to perturbation in the values of `ATPconsumption_Vm, HEX1_Vm, IRS_Katp_ratio, IRS_hillKatp_ratio`. The vertical line indicates the model value.

### 2.6 Sensitivity analysis of parameters affecting GSIS

To determine how the model parameters affect the rate of insulin release, we performed a local sensitivity analysis (Sauro, 2020). Figure 8C shows the sensitivity of insulin flux to a 10% change in model parameter values at different glucose concentrations. The rate of insulin secretion depends on the ATP/ADP ratio, so perturbing parameters that affect ATP formation and consumption has strong effects. Figure 8D shows the highly sensitive parameters that have positive and negative effects on insulin secretion, including factors affecting ATP consumption, ATP formation, hexokinase, phosphofructokinase, and ATP/ADP-dependent insulin secretion.

In conclusion, our systematic pathway modeling workflow provides insights into the key mechanisms of GSIS in the pancreatic *β*-cell.

## 3 DISCUSSION

We have developed a comprehensive kinetic model of GSIS in the pancreatic *β*-cell that can simulate glucose-dependent changes in glycolytic intermediates, ATP/ADP ratio, and their effect on insulin secretion. The main objective of this study was to establish a standardized workflow for data integration and normalization to construct a tissue-specific model of glycolysis and GSIS in the *β*-cell. Although we did not model other important pathways related to ATP homeostasis, such as the citric acid cycle, the pentose phosphate pathway, and the respiratory chain, our workflow can be easily extended to include them. Incorporating these pathways into our model will enable us to explicitly model the regulatory effect of downstream metabolites on the ATP/ADP ratio and insulin secretion. Previous studies have shown that fatty acids and amino acids can also induce insulin secretion in addition to glucose. Therefore, linking glucose metabolism with fatty acid and amino acid metabolism could help in understanding the insulinotropic effects of other fuel sources.

The increase in ATP levels triggers a cascade of events that culminate in the release of insulin from *β*-cells. Precisely, high ATP levels prompt the closure of ATP-sensitive potassium channels (Ashcroft, 2006). Consequently, the cell membrane depolarizes, opening voltage-gated calcium channels, which allows calcium influx. The influx of calcium triggers exocytosis of insulin-containing vesicles, leading to the release of insulin into the bloodstream (Rorsman and Braun, 2013; Guerrero-Hernandez and Verkhratsky, 2014). These electrophysiological changes resulting in insulin secretion were not modeled explicitly, but the effect of the ATP/ADP ratio on insulin secretion was modeled using a phenomenological (Hill-type) expression. Consequently, the model’s predictive capacity is limited to the steady-state glucose-insulin secretion dynamics. Expanding the model to explicitly describe these phenomena would allow to study experimentally observed patterns such as biphasic insulin secretion (Pedersen et al., 2008). Of note, the dynamics changing glycolytic intermediates were correctly described by the model.

Although our model has some limitations, it represents the first data-driven approach to integrate information from diverse sources and experimental setups. Moreover, it provides the first systematic analysis of the glycolytic changes that occur during insulin secretion in response to different glucose levels. Our study reveals that in GSIS, almost all glycolytic intermediates increase in a glucose-dependent manner as do total ATP and NADH, which is a significant finding.

Our model was developed to address the limitations of existing pancreatic *β*-cell models of glucose-insulin kinetics. These models often suffer from several drawbacks such as limited evaluation to a single data set, non-standardized formats of experimental data and kinetic parameters, and non-reproducible formats. To overcome these limitations, we have created open, free, and FAIR assets that can be used for the study of pancreatic physiology and GSIS. These assets include a fully reproducible SBML model of pancreatic *β*-cell glycolysis, a data curation workflow, strategies for unit and data normalization, and a large database of metabolic data of the pancreatic *β*-cell. Our systematic model-building workflow can be used as a blueprint to construct reproducible kinetic models of cell metabolism.

Computational modeling faces a significant challenge due to the substantial variation in data across different experimental systems, species, and cell lines. Often, relative data instead of absolute data is reported, further complicating the task of data integration. In this study, we developed a reliable data normalization workflow that was applied to experimental and clinical data from 39 studies conducted over the past 50 years on pancreatic, islet, and *β*-cell function in various species and cell lines. Our approach substantially reduced data heterogeneity and revealed a highly consistent response in glycolytic metabolites and insulin secretion. The high degree of conservation in the system of GSIS may have contributed to the effectiveness of the normalization workflow, as similar mechanisms are at play in different species, and the general changes can be observed across various experimental systems.

The study has laid a strong groundwork for enhancing our comprehension of the underlying reasons behind impaired insulin secretion. By mapping proteomics or transcriptomics data onto specific pathways, the developed model could be utilized to gain further insight into changes in GSIS, for instance in diabetic patients.

Furthermore, this model can serve as a crucial component for physiological whole-body models of glucose homeostasis, allowing researchers to investigate the relationship between insulin release and glucose uptake by insulin-responsive tissues.

In conclusion, this study utilized a systematic modeling workflow to gain insight into the key mechanisms involved in glucose-stimulated insulin secretion (GSIS) in pancreatic *β*-cells. When extended for translational purposes in clinical settings, it can serve to create reference models to identify variations in subjects which can lead to useful inferences regarding underlying metabolic conditions with therapeutic relevance.

## 4 METHODOLOGY

The workflow for building the kinetic model is illustrated in Fig. 2, with the following sections providing information on the individual steps.

### 4.1 Stoichiometric model

Chemical formulas and charges were assigned to all metabolites, and reactions were examined to ensure that they maintained mass and charge balance. The kinetic model encompasses glycolytic reactions and correlates the energy status of the *β*-cell with insulin secretion. sbmlutils (König, 2022c) was used to create and validate the model, while cy3sbml (König et al., 2012b) was used to confirm its coherence. The mass and charge balance of the system was verified using cobrapy (Ebrahim et al., 2023).

### 4.2 Metadata integration

Adding semantic annotations to models is an essential aspect of improving their interoperability and reusability, as well as facilitating data integration for model validation and parameterization (Neal et al., 2019, 2020). To describe the biological and computational significance of models and data in a machine-readable format, semantic annotations are encoded as links to knowledge resource terms. Open modeling and exchange (OMEX) metadata specifications were employed to annotate model compartments, species, and reactions with metadata information (Fig. 2B).

#### Case study: Phosphoglycerate kinase

The enzyme phosphoglycerate kinase (*PGK*) catalyses the conversion of 1,3-biphosphoglycerate (*bpg13*) and ADP to form 3-phosphoglycerate (*pg3*) and ATP.

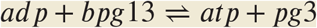

In our model, `PGK` is described by the following annotations:

SBO:0000176, vmhreaction/PGK, bigg.reaction/PGK, kegg.reaction/R01512, ec-code/2.7.2.3, biocyc/META:PHOSGLYPHOS-RXN, uniprot:P00558, uniprot:P07205.

The model components, including physical volumes, reactions, metabolites, and kinetic-rate laws, were annotated using Systems Biology Ontology (SBO) terms, which describe the computational or biological meaning of the model and data (Courtot et al., 2011). Biomedical ontology services such as Ontology Lookup Service (OLS) (Cote et al., 2010), VMH (Noronha et al., 2019), and BiGG (King et al., 2016) were used to collect these terms. Additional information for species and reactions were gathered from various databases such as HMDB, BioCyc, MetaNetX, ChEBI, and SEED. For instance, the model’s metabolites were annotated with identifiers from VMH, BiGG, KEGG, HMDB, BioCyc, ChEBI, MetaNetX, and SEED, while reactions were annotated with VMH, Rhea, MetaNetX, SEED, BiGG, BioCyc, and KEGG identifiers (Hari and Lobo, 2022). Enzymes catalyzing reactions were annotated with identifiers from enzyme commission (EC) numbers, UniProt (The UniProt Consortium, 2017), and KEGG. Finally, the annotations were incorporated into the SBML file using sbmlutils (König, 2022c) and pymetadata (König, 2022b).

#### Case study: 1,3-biphosphoglycerate

There is currently a bottleneck in data integration due to the use of multiple synonyms to refer to a single compound in data repositories. For instance, `bpg13` is identified by different names in SABIO-RK (*Glycerate 1,3-bisphosphate, 3-phospho-D-glyceroyl phosphate*) and BRENDA (*3-phospho-D-glyceroyl phosphate*). Additionally, the labeling of *1,3-biphosphoglycerate*, abbreviated as *DPG*, varies across existing *β*-cell models (e.g., *1,3-bisphospho-D-glycerate* in (Jiang et al., 2007) and *1,3-biphosphoglycerate* in (Salvucci et al., 2013)). Overall, `bpg13` is associated with seven synonyms: *1,3-Bisphospho-D-glycerate, 13dpg, 3-Phospho-D-glyceroylphosphate, Glycerate 1,3-bisphosphate, 3-phospho-d-glyceroyl-phosphate, 1,3-diphosphoglyceric acid, 3-Phospho-D-glyceroyl phosphate*. This issue makes it difficult to integrate data and information from different resources, highlighting the need to link chemical entities in the model to knowledge resource terms.

In our model, `bpg13` is clearly described by the following metadata annotations: SBO:0000247, vmhmetabolite/13dpg, bigg.metabolite/13dpg, kegg.compound/C00236, biocyc/META:DPG, CHEBI:16001, inchikey:LJQLQCAXBUHEAZ-UWTATZPHSA-N.

The formula and charge of`bpg13` are C3H4O10P2 and -4, respectively.

### 4.3 Kinetic parameters

Kinetic parameters, such as half-saturation constants (*K*_*M*_), inhibition constants (*K*_*I*_), activation constants (*K*_*A*_), and equilibrium constants (*K*_*eq*_), were gathered from literature and a variety of databases (see Fig. 2C). Values were programmatically accessed from UniProt (The UniProt Consortium, 2017), BRENDA (Placzek et al., 2017) using brendapy (König, 2022a), and SABIO-RK (Wittig et al., 2018). These databases were searched using an organism’s NCBI taxonomy identifier and reaction EC number as input search terms. Various parameters, including measurement type (*K*_*m*_, *K*_*i*_, and *K*_*a*_), experimental conditions (pH, temperature), KEGG reaction identifiers, enzyme type (wildtype or mutant), associated metabolite identifiers (SABIO compound name or BRENDA ligand id), UNIPROT identifiers associated with the isoforms of an enzyme, source tissue, and details of data source (PubMed identifier) were obtained. Since there is limited availability of kinetic data for *Homo sapiens*, we also searched for parameter values reported in studies of animal species that are closely related to humans and utilized them if no data were available for humans.

### 4.4 Synonym mapping

To map compound synonyms associated with each queried metabolite, we utilized compound identifier mapping services and available metadata annotations. First, we associated the name of each compound with internal database identifiers, such as the internal identifier of Glycerone-phosphate in SABIO, which is 28. Then, we linked the internal identifiers to external identifiers, such as those from ChEBI and KEGG. The external identifiers associated with the SABIO ligand identifier were obtained from cross-ontology mappings available in SABIO-RK. Similarly, we queried the REST API of UniChem to obtain the external identifiers associated with the BRENDA ligand identifier. By doing so, we were able to map most of the kinetic parameters to their respective compounds (Fig. 2D).

### 4.5 Model parameters

For each parameter in the model, the median value was calculated after synonym mapping and the values were assigned to the model parameters, see Fig. 2E. This was performed for initial concentrations, equilibrium *K*_*eq*_ constants, half-saturation constants *K*_*m*_, inhibition *K*_*i*_, and activation *K*_*a*_ constants.

### 4.6 Data curation

The next step involved curating data from studies that reported metabolite values, insulin secretion, or maximal velocities of glycolytic reactions *V*_*max*_ in pancreatic, islet, and *β*-cell lines (Fig. 2F). Relevant studies were identified through a literature search in PubMed, with a focus on time course and dose-response profiles of metabolite concentrations for metabolites and insulin secretion. Tissue homogenates were prepared by isolating islets from rodents, humans, or insulin-secreting cell lines (see Tab. 1). Assays were performed by stimulating the medium with various pre-incubation and incubation concentrations of glucose. To curate the data, established curation workflows from PK-DB (Grzegorzewski et al., 2021b), which were applied in a recent meta-analysis (Grzegorzewski et al., 2021a), were used. The numerical data was digitized by extracting the data points from the figures and tables using WebPlotDigitizer (Rohatgi, 2021). The incubation time and glucose concentration of the stimulation medium were recorded for all measurements, and meta-information such as organism and tissue type were documented.

The data is available under a CC-BY 4.0 license from https://github.com/matthiaskoenig/pancreas-model. In this study, version 0.9.5 of the data set is used (Deepa Maheshvare and König, 2023).

### 4.7 Unit normalization

The data measured in different studies is often reported in different units. Therefore, unit normalization was performed to integrate the data and convert metabolite concentrations and insulin secretion to standardized units of mmole/l (mM) and nmole/min/ml (*β*-cell volume), respectively (Fig. 2G).

Absolute measurements reported in metabolic profiling studies were found in various units such as per gram DNA, per gram wet weight or dry weight of the islet tissue, per cell, per islet, etc. To use these values for model calibration, both the absolute and relative measurements were first converted to concentration units in mM. The absolute values were converted to model units by multiplying the raw values with appropriate unit conversion factors. For instance, the islet content of glucose 6-phosphate, G6P, (pmol/islet) was converted to concentration units (mM) using the distribution volume of water in the islet (2nl/islet) (Ashcroft et al., 1970) as the conversion factor. Relative measurements were mainly reported with reference to a basal concentration. These relative measurements were converted to absolute quantity by multiplying the fold values with the respective metabolite concentration at the basal or pre-incubation concentration of glucose.

### 4.8 Data normalization and integration

Data collected from experiments performed in different laboratories, under different experimental conditions, and with different animal species showed significant variability after unit normalization. Therefore, data normalization was performed to eliminate systematic discrepancies between data reported in different studies (as shown in Fig. 2H). To achieve this, least squares optimization was used to minimize the distance between individual experimental curves and the weighted average of all curves for a given metabolite. The data normalization process involved a two-step procedure in which the steady-state data were first normalized for each metabolite. The resulting steady-state normalization was then used to normalize the time course data for that metabolite (see Fig. 4 for the example of glucose-6 phosphate).

#### 4.8.1 Steady-state data normalization

Steady-state (ss) experiments consisted of pre-incubation with one glucose dose followed by incubation with another glucose dose. The steady state data of the experiment *α*, 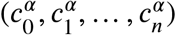 observed at *n* incubation glucose doses 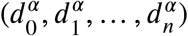 is expressed by the piecewise linear-interpolation function *C*^*ss*^. Here, *α* belongs to the set of steady-state experiments 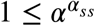 with 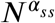 being the number of steady-state experimental curves of the metabolite *s*.

##### Mean curve

The mean steady-state curve 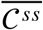 of each metabolite *s* is calculated as the weighted average of all experimental curves. The data points of the mean curve were interpolated using a piecewise smooth spline function. For data sets consisting of 2 data points, a linear interpolation was used.

We formulate a least-squares optimization problem to minimize the distance between the individual experimental curves and the mean curve 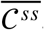. The cost function **F** of the optimization problem is given by,

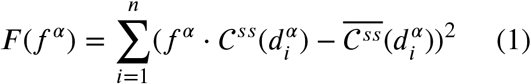

In Eq. 1, 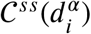 and 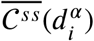 are the function values of the individual and mean interpolation function at the *i*^*th*^ value of the glucose dose. N is the number of glucose values in the dose-response curve of the experiment *α*.

For each experimental curve, the factor *f* ^*α*^ was determined so that the residual error in Eq. 1 is minimized. The residual error is minimum at the point where the derivative of the cost function **F** is zero. Taking the partial derivative of Eq. 1 with respect to the scale transformation parameter gives factor *f* ^*α*^ of the experimental curve *α* (Eq. 2).

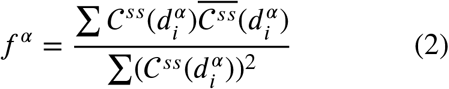

The scale factors of all steady state curves 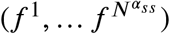 were determined by minimizing the respective cost functions 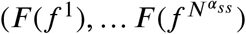. Multiplying the experimental curve *C*^*α*^ by the scaling factor *f* ^*α*^ shifts the experimental curve towards the mean curve. A new mean curve can be calculated with the scaled data. The curves were scaled iteratively until all *f* ^*α*^ converged.

#### 4.8.2 Time course data normalization

Time course (tc) experiments consisted of pre-incubation with one glucose dose followed by incubation with another glucose dose. The time-dependent data of the time course experiment 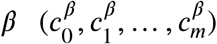 observed at *m* time points 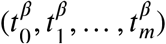 is expressed by the piecewise linear-interpolation function *C*^*β*^. Here, *β* belongs to the set of time course experiments 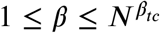 with 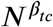 being the number of time course experimental curves of the metabolite *s*. For normalization, each time course was scaled by a factor *f* ^*β*^.

For a given incubation glucose dose *d*^*β*^, the metabolite concentration at the last time point 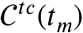 corresponds to the steady state value reached for the given *d*^*β*^

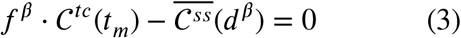

The scaling factor for the time course experiment follows as

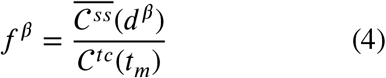

### 4.9 Model inputs

The SBML model was generated by specifying initial concentrations, rate expressions, parameter values, and compartmental volumes as the model inputs, see Fig. 2I.

#### Volume

The physical volume of the cytoplasmic compartment and the *β*-cell volume were obtained from the values reported in a morphometric study of *β*-cells (Dean, 1973).

#### Initial concentrations

The initial concentrations of glycolytic intermediates were obtained from the mean curve 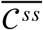 (Sec. 2.1) at a basal glucose concentration of 3 mM. The initial value of glucose in the external/blood compartment is 3 mM.

The initial concentrations of cofactors were expressed as polynomial functions passing through the data points of the mean curve, which is computed as the weighted average of data normalized experimental curves (Sec. 2.1). In the SBML model, the polynomial expressions were defined using assignment rules.

#### Kinetic constants

The median values of the half-saturation or Michaelis-Menten constants *K*_*m*_ (Sec. 4.5), were assigned to the model parameters.

#### Equilibrium constants

The values of the equilibrium constants *K*_*eq*_ were collected from NIST (Goldberg and Tewari, 2003) and EQUILIBRATOR (Noor et al., 2013).

#### Model equations

For all the glycolytic reactions, the biochemical interactions were expressed using modular rate laws (Liebermeister et al., 2010) of the form Eq. 5.

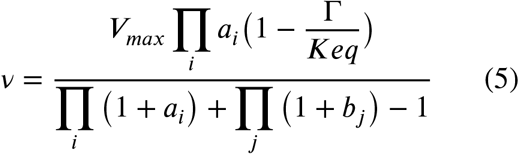

Here, *a*_*i*_ is *S*_*i*_/*Km*_*s*_, *b*_*i*_ is *P*_*i*_/*Km*_*p*_, S refers to the substrate and P refers to the product. *K*_*eq*_ is the equilibrium cons ant and Γ is he mass-action ratio (Liebermeister et al., 2010).

The use of detailed mechanistic rate laws was avoided due to the challenges associated with finding a large number of parameter values.

Insulin secretion was modeled via a phenomenological equation depending on ATP/ADP ratio. The insulin release flux given by Eq. 6, is characterized by three parameters, the maximal rate of insulin release *V*_*max*_, the Hill coefficient `n`, and *K*_*m*_ the ratio of ATP/ADP that results in half-maximal insulin release.

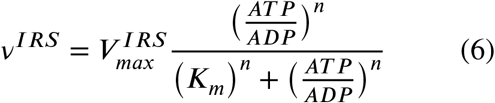

#### Boundary metabolites and reactions

Species in the external and mitochondrial compartments were assumed to be boundary species with constant concentrations, i.e. glucose and lactate in the external compartment and pyruvate in the mitochondrial compartment were held constant. Some boundary reactions were modeled as irreversible reactions, i.e. the export of lactate and the transport of pyruvate in the mitochondrion.

#### Metabolites determined by rate rules

To account for glucose-dependent changes in the concentrations of phosphate, NAD, and NADH, polynomial functions were used to express the concentrations as rate rules. This approach ensured that the concentration of fixed metabolites in the system increased as a function of glucose dose.

#### Changes in total adenine nucleotides

The sum of adenine nucleotides (*AT P* + *ADP* = *AT P*_*tot*_) changes with glucose. To account for these changes, a reaction Δ`ATP` was added that changes the total ΔATP according to the observed steady-state data for a given glucose value (Eq. 7).

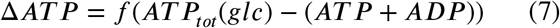

The *AT P*_*tot*_(*glc*) values are determined by the interpolating polynomial of the mean steady-state glucose dose response of the ATP+ADP data.

### 4.10 Model calibration

The normalized time-course and steady-state data was used for model calibration and parameter estimation (Fig. 2J). An overview of the subset of data used for model calibration is shown in Fig. 1. The following data were not used: NADH and NAD were fixed metabolites in the model, with NAD/NADH and NADH+NAD calculated from the metabolites. Total ATP was calculated by summing ATP and ADP, and ATP ratio was calculated by finding the ratio. The insulin secretion rate (IRS) was used to derive the parameters of the IRS function.

A subset of the *V*_*max*_ parameters was optimized to minimize the error between model predictions and experimental observations. The cost function is given by the sum of squares of residuals

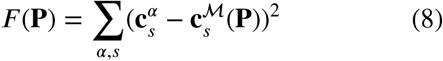

In Eq. 8, 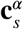 is the concentration of the metabolite *s* in the experiment *α* and 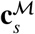 is the concentration of the metabolite *s* predicted^*s*^ by the model ℳ. **P** is the set of 16 parameters of maximum reaction rates *V*_*max*_. The experimental data of all transient metabolites in the model were stored in spreadsheets. The parameter estimation simulation experiments were set up using basiCO (Bergmann, 2023), the Python interface of COPASI (Hoops et al., 2006). The incubation glucose concentration and incubation time were mapped to the independent variable (*glc*_*ext*_, glucose in the external compartment) and model time, respectively. The transient metabolites were assigned to the model elements as dependent variables. The mean values of *V*_*max*_ calculated from the curated values of the enzyme activities were assigned as initial values. The lower and upper bounds specified for the reaction rates *V*_*max*_ were set to 0 and 10000, respectively. The calculations were performed using Cloud-COPASI, the front-end to a computer cluster at the Centre for Cell Analysis and Modelling. Cloud-COPASI is an extension of Condor-COPASI (Kent et al., 2012). 400 iterations of parameter estimation were performed on Cloud-COPASI using the SRES algorithm, a global optimization method. The optimal values of the parameter set were obtained from the iteration that yielded the minimum objective value and updated in the model.

### 4.11 Kinetic model and model predictions

All information was written into the model, validation was performed using sbmlutils, and model simulations were performed, see Fig. 2K, L.

Finally, we performed model predictions of glycolytic intermediates and insulin response as a function of varying glucose concentrations. The set of differential equations was numerically integrated using basiCO (Bergmann, 2023) based on COPASI (Hoops et al., 2006) and sbmlsim (König, 2021) based on libroadrunner (Welsh et al., 2023; Somogyi et al., 2015). For the glucose dose-response, glucose was varied as `linspace(0.01, 35, num=11)` and the model was simulated to steady-state. For the time course simulations, glucose was varied identically and simulations were run for 60 min. Simulations were performed either with COPASI or independently using libroadrunner to ensure reproducibility of key model results.

The model is available in SBML (Hucka et al., 2019; Keating et al., 2020) under a CC-BY 4.0 license from https://github.com/matthiaskoenig/pancreas-model. In this study, version 0.9.5 of the model is presented (Deepa Maheshvare and König, 2023).

## AUTHOR CONTRIBUTIONS

DM, SR, MK, and DP conceived and designed the study. DM and MK developed and implemented the computational model and data normalization workflow, and performed the analysis. DM curated the experimental data, performed parameter estimation, and drafted the initial version of the manuscript. All authors read, discussed the results, revised, and approved the manuscript.

## ACKNOWLEDGMENTS

DM thanks Dr. Murthy Madiraju S.R., Montreal Diabetes Research Center, CRCHUM, Montréal, Canada, Dr. Pedro Mendes, University of Connecticut, and Dr. Frank Bergmann, University of Heidelberg for the invaluable discussions. Access to Cloud-COPASI is supported by NIH Grant R24 GM137787 from the National Institute for General Medical Sciences.

